# Evolutionary exploration of drug-like chemical space utilizing generative AI and virtual screening

**DOI:** 10.64898/2026.03.26.714527

**Authors:** Christopher Secker, Philipp Secker, Fatih Yergöz, M. Özgür Celik, Surahit Chewle, Maria Phuong Nga Lê, Mustafa Masoud, Steffen Christgau, Marcus Weber, Christoph Gorgulla, AkshatKumar Nigam, Robert Pollice, Christof Schütte, Konstantin Fackeldey

## Abstract

The identification of suitable lead molecules in the vast chemical space is a critical and challenging task in drug discovery campaigns. Recently, it has been demonstrated that large-scale virtual screening provides a powerful approach to accelerate the identification of novel drug candidates by screening ever increasing virtual ligand libraries, which have reached magnitudes of *>* 10^20^ compounds. However, this desirable increase in potentially bioactive molecules poses a new challenge as enumerating and virtually screening such huge compound libraries is computationally prohibitive. Consequently, advanced approaches to navigate ultra-large chemical spaces and to identify suitable candidate molecules therein are urgently needed. Here, we present an evolutionary algorithm framework using molecular generative AI, reaction-based substructure searching, and iterative model fine-tuning for a targeted and efficient exploration of chemical fragment spaces. Combining this approach with large-scale virtual screening we are able to identify target-specific candidate molecules within the commercially available Enamine REAL Space (∼10^15^). We demonstrate the applicability of the approach by successfully identifying and biochemically validating pH-specific ligands of the *µ*-opioid receptor. Our results demonstrate that integrating generative AI with evolutionary algorithms provides a promising route to explore ultra-large chemical spaces for the discovery of novel, synthetically accessible lead molecules.

## Introduction

The drug-like chemical space encompasses approximately 10^60^ small molecule compounds [1, 2]. Many virtual chemical libraries reflecting parts of this vast chemical space have been established to date. One of them, the Enamine REAL Space, expanded from initially 3.5 million to over 70 billion synthetically accessible molecules in 2016 [3]. As of today, the extended Enamine REAL Space comprises trillions of “make-on-demand” compounds [4]. These numbers stand in big contrast to the physical chemical libraries typically used in high-throughput screening (HTS). Such libraries usually consist of up to 1–5 million compounds and therefore often lack representation of various chemical classes and scaffolds [5]. Consequently, the much larger virtual libraries are likely to contain (1) more potent hit molecules and (2) more structurally diverse hit molecules applicable for drug discovery campaigns compared to the compound collections used in traditional HTS.

Structure-based virtual screening represents one efficient technique to search for potential new lead candidates within such large virtual chemical libraries [6]. Virtual hits are usually identified by computing or approximating, for each candidate molecule, its change in Gibbs free energy upon binding (binding free energy difference, Δ*G_b_*). As processing resources and techniques have advanced, the concept of “ultra-large” virtual screening has evolved, with the number of molecules to be individually screened reaching into the billions [7]. Recent advances in high-performance computing, improved CPUs and GPUs, parallel computing, and cloud computing have significantly augmented our capacity to efficiently orchestrate and process such extensive tasks [8, 9]. Additionally, machine learning (ML)-based methods yield techniques for managing such ultra-large virtual molecule libraries and potentially conducting guided searches therein more efficiently.

In a complementary approach, generative artificial intelligence (AI)-based molecular models have recently been shown to enable the de novo design of molecules for *in silico* drug discovery, instead of screening ultra-large libraries [10–12]. Especially, the combination of physics-based and empirical methods for scoring ligands (e.g., free energy perturbations or molecular docking) with generative AI models has proven potential to identify novel high-affinity binders [11, 13]. However, one major challenge of this approach is to ensure that the AI-generated molecules are physically tangible, i.e., that they can be chemically synthesized within reasonable time frames and at efficient costs. This is especially important when using rather fast but coarse scoring approaches, such as molecular docking, where relatively low experimental hit rates must be expected [13–15]. Therefore, it is essential that the virtually selected hits are readily synthesizable to allow for fast experimental evaluation. The two complementary approaches, virtual screening of ultra-large ligand libraries and generative AI-based ligand design, can potentially be combined in evolutionary workflows to efficiently yield novel and, at the same time, easily tangible hit molecules for drug discovery campaigns.

Evolutionary based algorithms have previously been shown to provide a systematic framework to efficiently search vast chemical space [16–18]. They are inspired by evolutionary processes in nature and are part of stochastic heuristic optimization methods. In an *in silico* drug discovery workflow, such algorithms, e.g., start with a first assessment of a starting population’s predicted binding potential based on, e.g., docking scores (Fig 1.1). Iteratively, a selection of the best performing individuals is followed by variation through recombination and mutation. The newly formed population (*successor generation*) is then evaluated, and a new selection and variation cycle is initiated. However, it is well known that evolutionary algorithms cause populations to converge too quickly, leading to the discovery of only local optimal solutions (*premature convergence*) [19]. Another drawback can be the limited diversity within the population, which ultimately results in only a small portion of the chemical space being explored [17]. These limitations can potentially be overcome by including AI-based molecular generative models for the recombination and mutation steps in an evolutionary workflow [20], which could help generate more diversified successor generations. Using generative molecular AI models for these crossover operations, in contrast to classical recombination and mutation approaches, could improve exploration and help identify target-specific global optima in chemical space. In this work, we present such an approach and applied it to the identification of novel pH-specific agonists of the *µ*-opioid receptor (MOR).

**Fig. 1.**
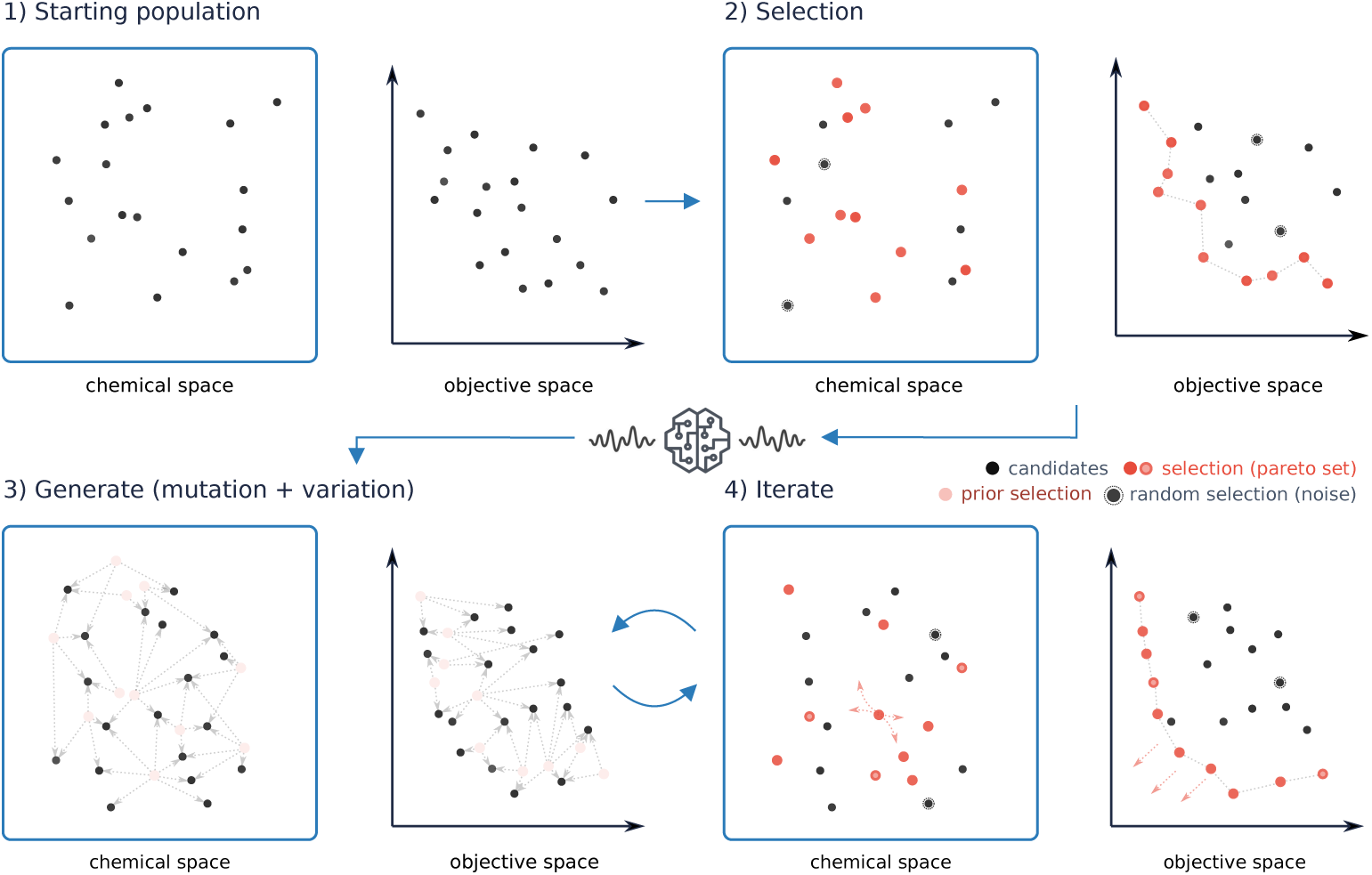
AI-directed ligand evolution for chemical space exploration. 1) Starting with a random set of molecules (“Starting population”), each individual (“candidates”, *dark gray data points*) of the population is scored and thereby transformed into an objective space reflecting, e.g., the predicted binding potential under different conditions. 2) The best performing individuals (“selection”, *orange data points*) are selected. In a multi-objective optimization scenario these represent, e.g., the Pareto set of molecules, i.e. the individuals of the population that meet best two conflicting conditions. 3) The selected best performing individuals serve as the training data used to fine-tune a pretrained generative model. The fine-tuned model is then used to generate a novel successor generation of candidate molecules (“mutation + variation”, *gray dotted arrows*). To improve diversity and counteract overfitting of the model, an additional random set of molecules (noise) is added to the training set. The novel population is again mapped into the object space (e.g. by molecular docking). 3) This procedure is iteratively repeated (“Iterate”) until a stop criterion is fulfilled (e.g. promotion of the Pareto front is below a defined threshold).

## Results

### Evolutionary and AI-guided exploration of chemical space

The main principle of the procedure conceived in this study can be outlined as follows: In the first step, a random starting population of molecules is generated (Fig. 1, *1) Starting population, chemical space*), and the binding potential of each individual in the population (*ligand*) to the target receptor(s) is assessed using molecular docking (Fig 1, *1) Starting population, objective space*). In the next step, the best performing individuals are selected, e.g. the Pareto-optimal ligands of the population. The Pareto set, e.g., consists of the molecules that have the best predicted binding potentials according to both criteria; e.g., the best compromise between a high predicted binding potential to a target receptor and a low predicted binding potential to, e.g., a risk-associated off-target (Fig 1, *2) Selection*. In the next step, a pretrained molecular generative model is fine-tuned with the best performing individuals of the first generation (Fig. 1, *selection*) and the fine-tuned model is then used to generate the subsequent offspring population of ligands. Additionally, a random set of molecules is added to the training set, which helps improve diversity within successor ligand populations (Fig. 1, *random selection*). The fine-tuning and sampling steps represent the generative components of the algorithm, responsible for updating and varying the information flow between generations and replacing the standard variation and mutation methods of classical evolutionary recombination approaches. The new generation of molecules thus emerges from, e.g., the top scoring (the *fittest*) or, in a multi-objective optimization scenario, also from the Pareto set of molecules (the best *compromise*) (Fig. 1, *3) Generate*. The new generation is again mapped to the objective space by, e.g., molecular docking. This process is iteratively repeated, through which the chemical space is explored in a way that the population improves with respect to the objective space (e.g., propagation of the Pareto front) (Fig. 1, *4*). The steps are repeated until the improvement of the population becomes insignificant. By iteratively fine-tuning the generative AI model over multiple subsequent iterations, the model not only performs a crossover operation based on the currently best performing individuals of the population but also memorizes features of the best individuals from multiple generations, thereby implicitly guiding the search in chemical space.

To provide efficient scalability of binding potential predictions, we combined this evolutionary workflow with the open-source, ultra-large virtual screening platform VirtualFlow [8, 21]. To conduct the generative AI-based variation and mutation step, we used one of the best performing and most efficient molecular generative models for chemical space exploration [22, 23]. To ensure that each generated molecule can be easily obtained for experimental validation, we additionally included a pipeline that automatically performs the decomposition of each AI-generated molecule [24] and verifies the availability of its substructures within the extended Enamine Real Space library of building block structures [4]. This approach can be viewed from two perspectives: (1) the matching of the fragmented AI-generated molecules with the building block library serves as a prefiltering step to enhance the chances of the synthesizability of the generated molecules, or in case of additional compliance with the reaction rules; (2) the molecular generative model is only able to suggest the next offspring population out of all possible building block combinations within the Enamine REAL Space. This strategy ensures that the workflow yields only synthetically accessible molecules, which is crucial for its practical applicability.

### Iterative optimization of pH-specific ligands

To apply and validate the presented pipeline, we conducted a search to identify pH-specific ligands of the MOR. Such substances have previously been shown to reduce the critical side effects of current opioid therapeutics [25]. For molecular docking, we first created a protein structure model of the human MOR based on a previously resolved x-ray crystallographic structure (PDB: 8EF5) [26]. The protein structure was initially resolved in the presence of fentanyl, thus representing an agonist-bound state of the receptor. For protein preparation, the structure was stripped of small molecular compounds, and side-chain protonation states were assigned at a neutral pH of 7.4 and an acidic pH of 6.5. The target area for molecular docking was centered on the bound fentanyl to prioritize compounds that address the same binding site. Interestingly, the prediction of the side chain protonation states at pH 7.4 and 6.5 revealed that, in the selected target region, the two protein structure models (pH 7.4 and 6.5) are predicted to have identical side chain protonation states. This is especially notable for Asp149, which was shown to engage in a key salt bridge interaction with the tertiary amine of the piperidine ring of fentanyl [26]. Due to the identical side chain protonation states and conformations at the target site of the two protein models, pH-specific ligands in our screening workflow are expected to primarily differ in the dominant protonation state of the ligands within their respective environments. One potential exception to this can be ligands that form contacts with Asp116, which is adjacent to the fentanyl binding site and is predicted to be deprotonated and negatively charged at neutral pH, but to be in a neutral, uncharged state at acidic pH. However, Asp116 is rather distant from the center of the target site and could therefore only be reached by larger, extended compounds.

For generating suitable candidate molecules in the workflow, we used an established molecular generative model that we first pretrained with ∼1 million compounds from the ChEMBL database (Fig. 2a, *Model pretraining*). To obtain the starting population, we sampled 1,000 random ligands from this pretrained model (Fig. 2a, *Molecule generation*), which then served as input for two separate pipelines of pH-dependent (pH 7.4 and 6.5) ligand preparation (Fig. 2a, *Ligand prep*). Each set of prepared ligands was used for docking against the respective MOR structure model prepared at pH 7.4 and 6.5 (Fig. 2a, *Docking*). After docking, the results were combined, and a total ranking of the compounds was generated (Fig. 2a, *Scoring & Ranking*). To prioritize candidate molecules that show higher potential binding predictions at acidic pH compared to neutral pH, this two-objective optimization task was transformed into a single-objective task in the form of optimizing (i.e., maximizing) a weighted average sum-based score (scalarization). By using random weights in scalarization, it was shown that the convex parts of the Pareto front can be approximated [27, 28]. To ensure equal contribution of each docking result, we first performed min-max scaling of the docking scores. Since a low binding free energy corresponds to a higher predicted binding potential, the min-max scaled score at the respective pH [*x^′^*(pH)] was inverted 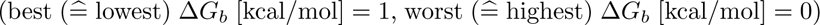:

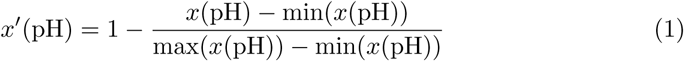

**Fig. 2.**
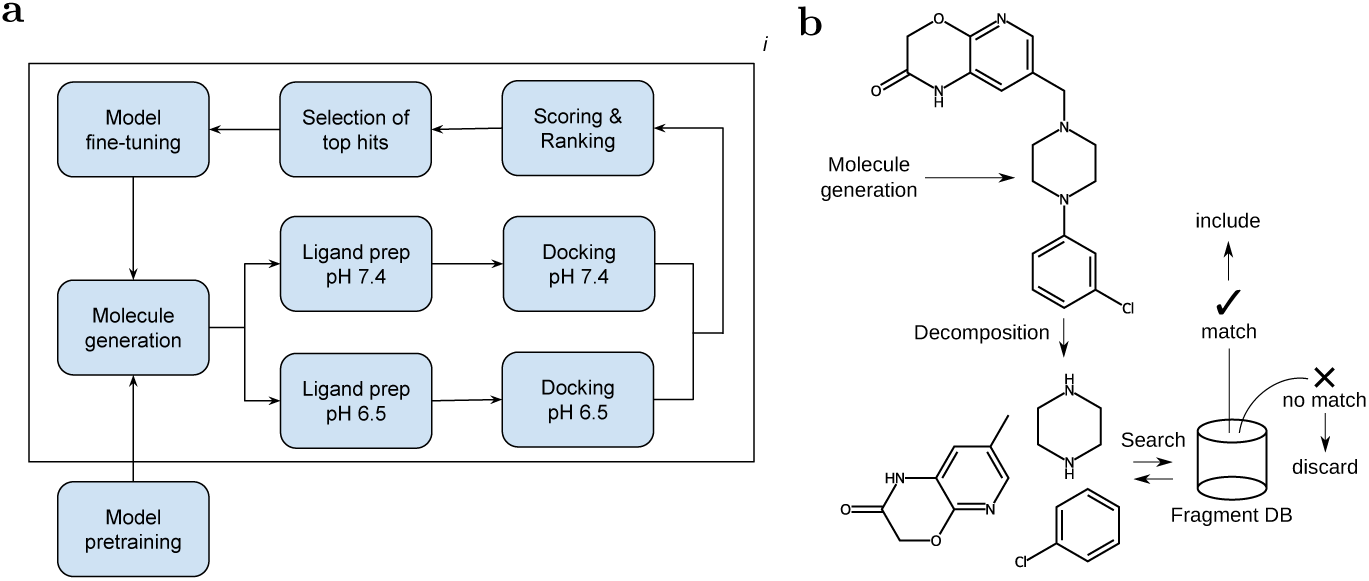
Generative AI-based virtual screening workflow to identify pH-specific ligands from fragment spaces. A pretrained molecular generative model is used to generate an initial, random set of ligands (*starting population*, 1,000 ligands). The candidate molecules are protonated at pH 7.4 and pH 6.5 and conformers are generated (Ligand prep). Prepared ligands are then virtually screened against the protein target protonated at pH 7.4 and pH 6.5, respectively. The results of the two docking scenarios are collected and scoring and ranking was performed (assessing *fitness*). The ligands ranking at the top 20% (*fittest*) are combined with ∼ 5% random ligands (*mutation*) used for fine-tuning the model (selection of individuals as *parents*). The fine-tuned model is then used to sample the next generation of ligands (*offspring*, 1,000 ligands). This process is iteratively repeated.

Weighted average sums for the docking scores at two different pH conditions [*x̄_w_*(pH_1_, pH_2_)] can then, in principle, be calculated as follows:

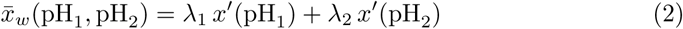

While using random weights with *λ*_1_*, λ*_2_ ∈ [0, 1] and *λ*_2_ = 1 − *λ*_1_ can lead to Pareto-optimal points, we have used fixed weights of 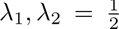 to prioritize compounds for which a high predicted binding potential at acidic pH contributes equally as a low predicted binding potential at neutral pH to the scalar product. Additionally, we inverted the scaled docking score at neutral pH [*x^′^*(pH_7.4_)] to prioritize ligands with a high predicted binding potential at acidic pH and a low predicted binding potential at neutral pH:

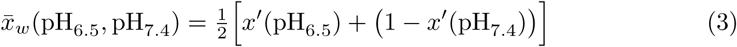

After each cycle of molecule generation, ligand preparation, and molecular docking, the ligands were ranked in descending order by *x̄^′^_w_* (pH_6.5_, pH_7_._4_), and the top 20% were used as training data (or *parents*) to refine the molecular generative model for generating the offspring in the next iteration. This loop of molecule generation, ligand preparation, docking, ranking, and model fine-tuning (Fig. 2a) was iteratively repeated for 20 iterations (or *generations*).

Analyzing the top ligands after each iteration, we observed that the mean average sum [*x̄^′^_w_* (pH_6.5_, pH_7.4_)] of the top 100 ligands increased over the iterations from ∼0.5160 ± 0.0006 at *i* = 1 to ∼0.5260 ± 0.0008 at *i* = 20, p *<* 0.0001 (Fig. 3a). An increase in the weighted average sum also led to molecules with slightly improved predicted binding affinities at acidic pH, as well as at neutral pH (Fig. 3b). However, there was a stronger improvement in the docking scores at acidic pH (∼-8.83 ± 0.07 kcal/mol at *i* = 20 for pH 7.4 and ∼-9.16 ± 0.06 kcal/mol at *i* = 20 for pH 6.5, p = 0.000433). Importantly, the mean difference between the estimated binding free energies [Δ*G_b_*(pH_6.5_) – Δ*G_b_*(pH_7.4_)] also showed an iteration-dependent increase, reaching a mean difference for the top 100 ligands generated in iteration step 20 of ∼0.33 ± 0.02 kcal/mol. Interestingly, the proportion of differentially protonated ligands among the top 100 of each iteration strongly increased over the course of the iterations. While in iteration step 1, only ∼5% of candidate molecules were predicted to be neutral at pH 7.4 but to contain a single proton at pH 6.5; after 20 iterations, ∼60% were predicted to be neutral at pH 7.4 but monoprotonated at pH 6.5 (Fig. 3d, pH 6.5 1H+). Due to the similarity of the side chain protonation states at the target site at the respective pH values, the ligand protonation state (Fig. 3d) is expected to be the main driver for differential binding potential predictions between the neutral and acidic environments (Fig. 3c,d). Accordingly, the workflow implicitly learns to generate and prioritize compounds that fulfill this crucial criterion.

**Fig. 3.**
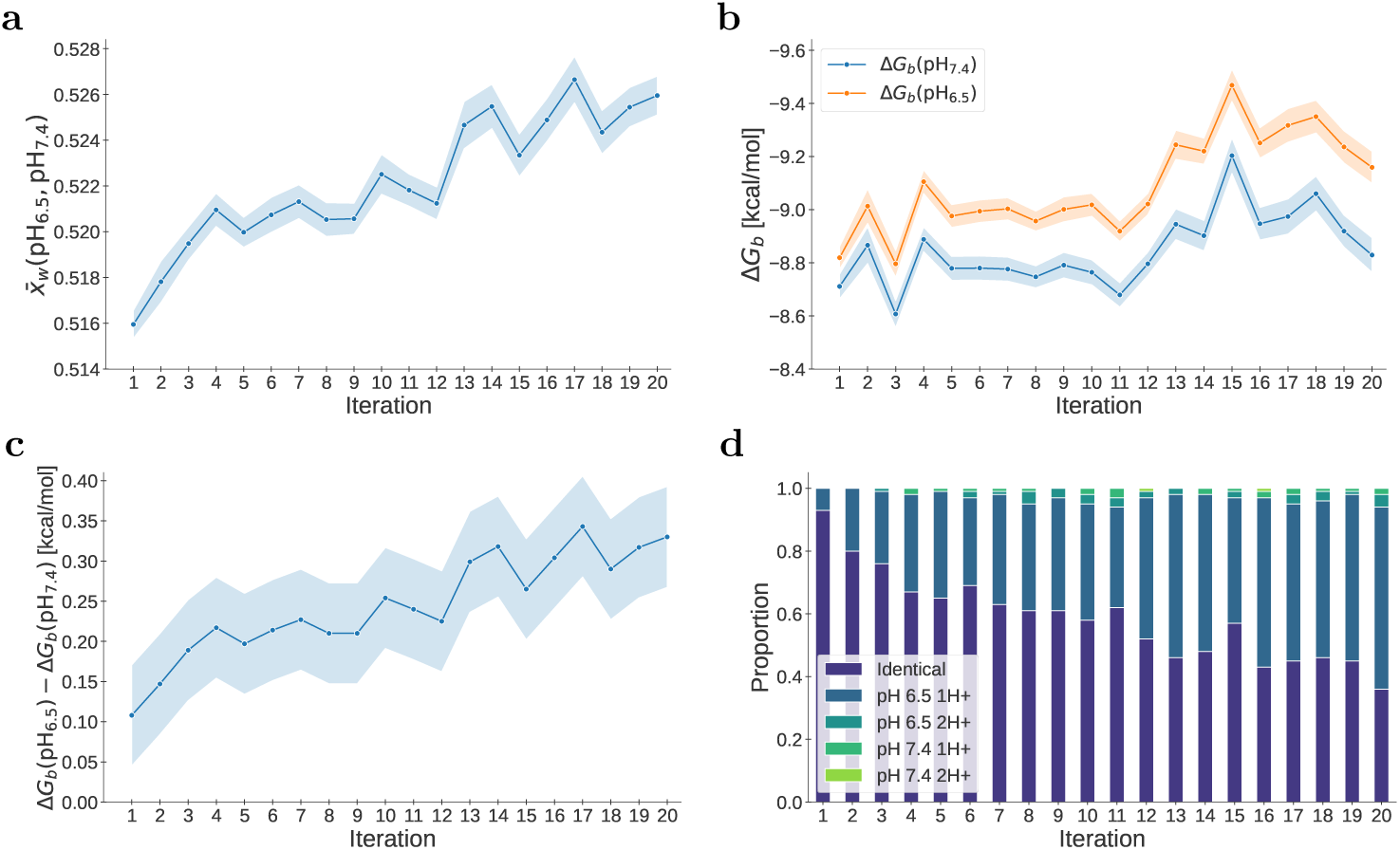
Optimizing for pH-specific MOR ligands. (**a**) Optimization of the objective function (weighted average sum, *x̄_w_*) over 20 iterations. The mean scores ± SEM of the top 100 ligands of each iteration, respectively, are shown. (**b**) Mean docking scores ± SEM (Δ*G_b_* [kcal/mol]) of top 100 ligands of each iteration at pH 7.4 and pH 6.5, respectively. (**c**) Mean difference of the docking scores ± SEM at the different conditions [Δ*G_b_*(pH_6.5_) - Δ*G_b_*(pH_7.4_)] of top 100 ligands of each iteration are shown. (**d**) Proportion of identical, monoprotonated (1H+) or diprotonated (2H+) ligands among the top 100 ligands of each iteration at pH 6.5 or pH 7.4, respectively.

### Selecting virtual hits for experimental validation

To select virtual screening hits for chemical synthesis and experimental validation, we chose ligands that showed a low binding free energy at acidic pH but a higher one at neutral pH. When plotting the total population of ligands from the workflow after 20 iterations (19,318 ligands), the most promising pH-specific candidates can be found in the lower right part of the graph. Interestingly, we could also observe that both binding free energies, at pH 7.4 (Fig. 4a, *green histogram*) and 6.5 (Fig. 4a, *yellow histogram*), increased over the course of the iterations. For further analyzes, we only considered compounds that exhibited a maximal binding free energy of −8 kcal/mol under either neutral or acidic conditions (Fig. 4a, *dotted lines*). We then selected the top 150 ligands from the ranking sorted by the weighted average score (3) in descending order (Fig. 4a, *orange data points*). To further prioritize the virtual hits, we next filtered for compounds that were predicted to be monoprotonated at pH 6.5, while they were predicted to be neutral at pH 7.4. Ligands with more than one protonation site were discarded since the micro-p*K_a_* prediction-based approach used for protonation state generation [29] was only implemented for ligands with a single ionizable site.

**Fig. 4.**
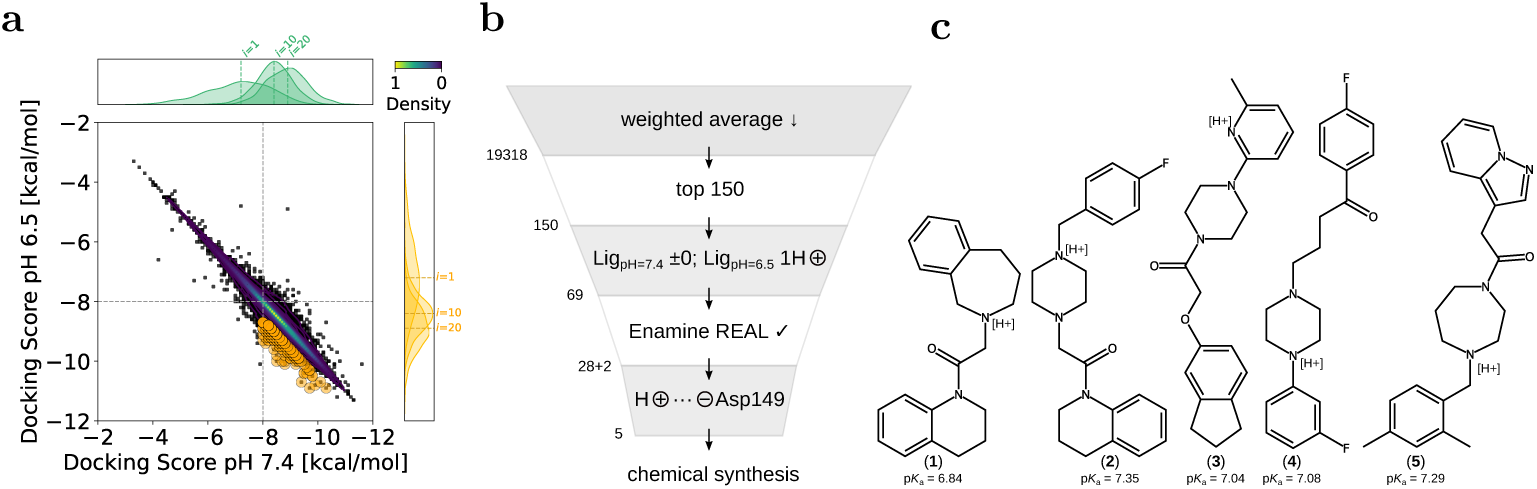
Selecting hits for experimental validation. Hit selection and synthesized candidate molecules. (**a**) Virtual screening results of all iterations (19,318 data points). The selected hits (top 150) are marked and consist of ligands that have a maximum weighted average score while scoring ≤ −8 kcal/mol at pH 7.4. (**b**) Hit prioritization. From the total 19,318 ligands, all were sorted in descending order by the weighted average score, then the top 150 were selected. Ligands were prioritized that are predicted to contain one positively charged hydrogen atom (H+) at pH 6.5, and are predicted to be neutral at pH 7.4. The remaining 69 ligands were evaluated by Enamine for synthesis compatibility within the REAL Space. Finally, hits from the Enamine REAL Space and two closely related hit analogs (28+2) were prioritized if they were predicted to form a salt bridge with Asp149 under acidic conditions, which would get lost when the respective ligand becomes deprotonated at neutral pH. Finally, five compounds were selected for chemical synthesis. (**c**) Two-dimensional structures of synthesized hit compounds. All compounds contain a nitrogen atom with a predicted acid dissociation constant of 6.5 *<* p*K_a_ <* 7.4, thus are predicted to be predominantly protonated and positively charged at pH 6.5 while being predominantly in a neutral state at pH 7.4.

Out of the top 150 ligands, 69 were predicted to be monoprotonated at pH 6.5 and neutral at pH 7.4. Of the 69 compounds, 28 were compatible for synthesis within the fragment and reaction space of Enamine REAL, while for two additional compounds, closely related analogs were available (+2). Finally, we manually inspected each of the remaining 30 ligands’ predicted binding modes (docking poses) and analyzed whether they are predicted to form a salt bridge between their positively charged hydrogen at pH 6.5 and the negatively charged Asp149 at the target site. This salt bridge represents a key contact for other MOR agonists, and the pH-dependent presence of the ligands’ hydrogen at this critical contact can result in strong pH-dependent binding [30]. Prioritizing structures that are predicted to form a salt bridge with Asp149 resulted in the selection of five compounds for synthesis (4b), which all contain one ionizable group in the range 6.5 *<* predicted p*K_a_ <* 7.4 (Fig. 4c). Interestingly, three of the compounds, similar to the potent MOR agonist fentanyl, contain a 1,4-diazine moiety (Fig. 4c, 2–4). All five compounds were successfully synthesized.

### Identification of a novel pH-specific MOR ligand

We next evaluated the pH-specific binding of the selected and synthesized hit compounds using a radio-labeled reporter displacement assay (RDA). This assay detects the displacement of radio-labeled [D-Ala^2^, N-MePhe^4^, Gly-ol]-enkephalin (DAMGO), a known peptidic MOR agonist, upon competitive binding. The binding assay was conducted under neutral as well as under acidic conditions, as previously described [25, 30]. All five synthesized compounds were tested for competitive binding from 10 nM to 50 µM. While one of the compounds did not show detectable displacement of radio-labeled DAMGO from the MOR, we observed a reduced radio-ligand signal in competitive binding experiments with compounds 1, 2, 4, and 5 (Supplementary Fig. 1). Among these, compounds 1, 4, and 5 demonstrated the anticipated pH-dependent activity profile, exhibiting enhanced inhibitory potency at pH 6.0 relative to pH 7.4 at equimolar concentrations. Compound 1 showed a clear concentration-dependent displacement of DAMGO at compound concentrations ≤ 1 µM (Fig. 5a, *compound 1*), but its effect was limited to displacing ∼20% of DAMGO. We further investigated the binding of compound 1 to the MOR and determined its half-maximal inhibitory concentration (IC_50_) under neutral and acidic conditions. Notably, compound 1 showed strong potency at acidic pH (IC_50_ = 167 ± 179 nM), while its potency was ∼10-fold lower in a neutral environment (IC_50_ = 1424 ± 541 nM) (Fig. 5b). Compound 4 generally exhibited lower potency but produced markedly stronger inhibition at higher concentrations compared to compound 1. At 50 µM, compound 4 reduces DAMGO binding by approximately 80% at pH 6.0 and about 40% at pH 7.4 (Supplementary Fig. 1). In summary, four out of five compounds exhibit at least moderate displacement of DAMGO, and three of these four compounds show increased activity at pH 6.0. Compound 1 demonstrates the highest potency; however, due to the low effect size, which is most likely caused by incomplete competition, the compound will require further optimization.

**Fig. 5.**
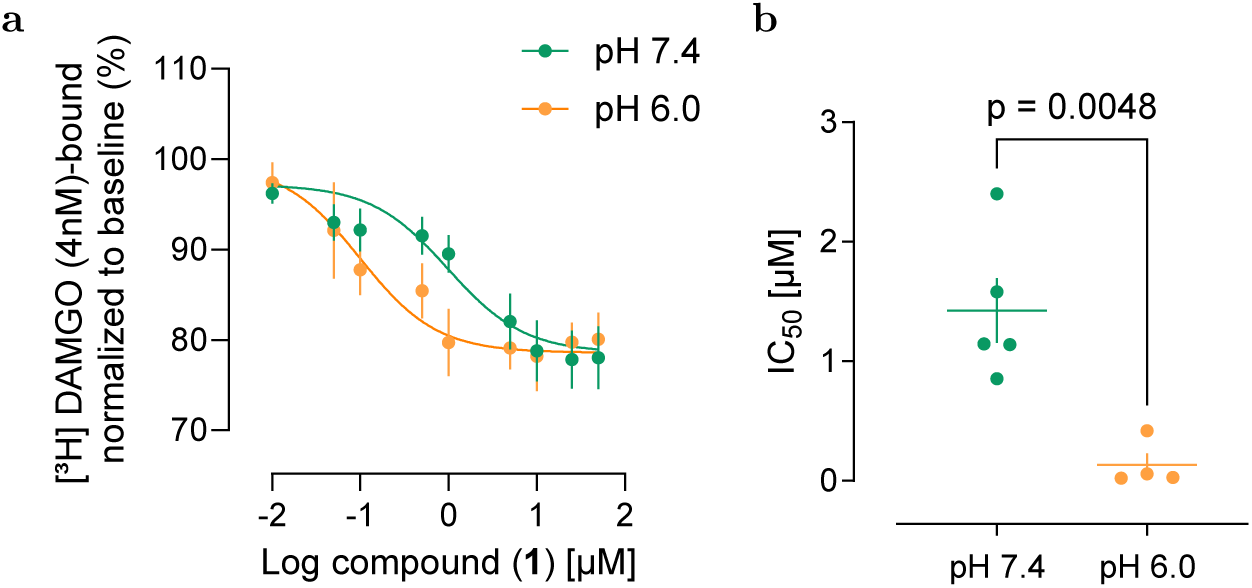
Experimental validation of pH-specific MOR ligand. (**a**) Radio-labeled RDA adding 10 nM to 50 µM of compound 1 to replace 4 nM DAMGO. Concentration series was conducted in a neutral (*green data points*, pH 7.4) and acidic (*orange data points*, pH 6.0) environment. Data points show mean ± SEM of minimum 4 independent experiments (*n* ≥ 4) (**b**) Comparison of half-maximal inhibitory concentrations (IC_50_) of compound 1 in a neutral and acidic environment. Horizontal lines indicate mean, vertical lines represent SEM. Compound 1 shows a significant higher potency at acidic (IC_50_ = 167 ± 179 nM) compared to neutral (IC_50_ = 1424 ± 541 nM) pH (two-tailed t-test, p = 0.0048).

To gain further insights into the compound’s mode of action, we additionally analyzed the binding mode and mechanism of its experimentally confirmed pH-specific binding to the MOR. Inspecting the predicted binding pose obtained from molecular docking, we found that, similar to e.g. fentanyl and morphine, it forms a strong salt bridge between its protonated heterocyclic nitrogen and the negatively charged Asp149 side chain. Additionally, it is predicted to form a *π*-anion contact with Asp149 and a *π*-*π* stacking interaction with the phenol ring of Tyr328. To investigate how deprotonation under neutral conditions affects these contacts and the predicted binding mode, we additionally performed molecular dynamics simulations of compound 1 in complex with the human MOR protein model. The trajectories obtained under neutral and acidic conditions revealed the differential strength of binding between the protonated and deprotonated compound 1. On the one hand, the protonated compound stays firmly inside the binding pocket, primarily through a salt bridge with Asp149. This stability in the binding pocket is further augmented by hydrophobic interactions with Ile146 and Met153, as well as *π*-*π* stacking interactions with Tyr328 (Fig. 6c, *left panel*). On the other hand, deprotonation of the nitrogen within the benzapine moiety breaks the salt bridge with Asp149, whereby the tetrahydroquinoline group of the deprotonated compound 1 gains conformational freedom, and the hydrophobic interactions with Ile143 dissipate along with the *π*-*π* stacking of Tyr328 (Fig. 6c, *right panel*), leaving the ligand largely unanchored.

**Fig. 6.**
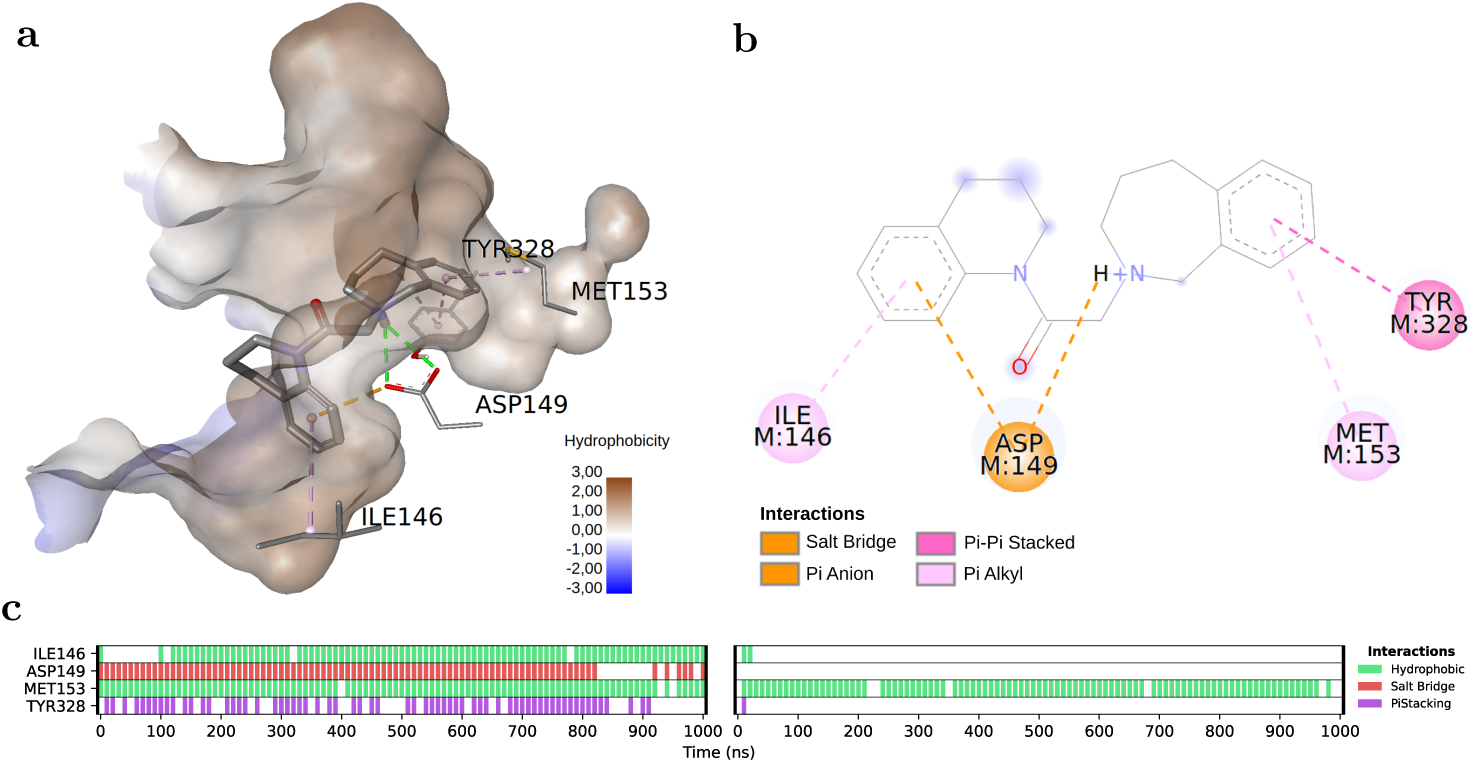
Protonated compound 1 is predicted to form key contacts at the fentanyl binding site. (**a**) Predicted binding mode of compound 1 at the target site. The protonated nitrogen is centered onto the negatively charged carboxylate of Asp149, while the aromatic ring of the benzazepine moiety is predicted to form a *π*-*π* stacking interaction with Tyr328. Displayed molecular surface is color-coded by hydrophobicity (*blue* indicating more hydrophobic, *red* indicating more hydrophilic). (**b**) Two-dimensional view of the ligand-side chain interactions at the predicted binding site and mode. The carboxylate moiety of Asp149 is predicted to form a salt bridge with the benzazepin-1-ium cation and a *π*-anion interaction with the 1,2,3,4-tetrahydroquinolin moiety of compound 1. The phenol ring of Tyr328 is predicted to engage in *π*-*π* stacking with the benzazepin moiety. Ile146 and Met153 are predicted to form *π*-alkyl interactions with the two *π* systems of compound 1. Particular attention was given to the salt bridge between the quaternary nitrogen on the ligand and the deprotonated oxygen of Asp149 as it was expected to strengthen the binding in low pH conditions. (**c**) Timeline of compound 1 contacts with the MOR during molecular dynamics (MD) simulations. Protonated compound 1 engages in strong interactions with the receptor over the simulation time of 1 µs (*left panel*). The salt bridge with Asp149 and *π*-*π* stacking with Tyr328 as well as hydrophobic interactions with Ile146 are lost as compound 1 is deprotonated (*right panel*).

## Discussion

Identifying novel lead molecules remains a central challenge in modern drug discovery campaigns. Hit molecules obtained from traditional HTS often require multiple iterative optimization cycles that are time-consuming, costly, and often have uncertain outcomes [31–33]. There is great hope that AI-based drug discovery can potentially help mitigate these challenges by, from the start, exploring a much larger chemical search space than physical libraries, thereby increasing the likelihood of identifying superior initial hits [34]. Efficiently navigating the immense and irregular chemical space while balancing multiple objectives, such as potency, synthetic feasibility, and drug-likeness, is uniquely complex. Since chemical space lacks a well-defined structure and gradient information, traditional optimization algorithms often struggle to explore it effectively [35, 36]. Evolutionary algorithms, with their stochastic and population-based nature, are particularly well suited for such problems, as they can iteratively refine populations without assuming a smooth or continuous search landscape [37].

Here, we introduce an evolutionary algorithmic framework that integrates molecular generative AI with high-throughput virtual screening to iteratively improve a population of candidate molecules for early-stage drug discovery. The presented workflow explicitly prioritizes synthetically accessible molecules by restricting generated candidates to a predefined set of precursor molecules (“building blocks”), thereby operating in so-called chemical “fragment spaces” [3, 38]. Although the approach does not guarantee strict adherence to known reaction rules, it substantially increases the probability that the proposed molecules can be synthesized through short and feasible synthetic routes. In a proof-of-concept study, we identified several novel candidate molecules from the ∼10^15^ large Enamine REAL Space [4], which showed a pH-specific predicted binding potential in the molecular docking readout. Notably, we experimentally confirmed the nanomolar, pH-dependent binding of one of the identified compounds to the MOR, demonstrating the practical applicability of the proposed approach for identifying potent ligands from large fragment spaces.

Other approaches to address the challenge of constantly growing ultra-large chemical libraries mainly pursue two differing strategies. One of them, utilized by tools like HIDDEN GEM [39], SpaceHASTEN [40] or Deep Docking [41], is to use similarity metrics or active learning models to predict top binders after docking a smaller fraction of the chemical library. While this approach can be computationally efficient, similarity-based methods usually require fully enumerated chemical libraries that demand extensive storage space and are cumbersome to extend when new molecules are added. While active learning-based methods can also use non-enumerated virtual libraries [40], their outcomes largely rely on the predictive capability of the applied models. Another approach circumvents the requirement of a fully enumerated library by operating on fragments or building blocks, which are connected through established chemical reactions to construct ultra-large libraries. Typically, the individual building blocks are docked first, and those with the best scores are subsequently combined with additional building blocks, thereby assembling the molecules step by step. The underlying hypothesis is that combining fragments with the highest binding scores will yield the best overall molecules. Such approaches include Chemical Space Docking [42] and V-SYNTHES [43]. While this stepwise procedure may be logical and efficient, it carries the risk of committing early to specific fragments, potentially resulting in limited chemical diversity and the loss of unexpected molecular structures whose binding potential could not be readily predicted from the individual fragments. Another drawback is that individual ligands emerge from multiple docking steps. Given that docking is inherently error-prone and imprecise, the likelihood of inaccuracies increases with the number of docking steps performed. Our approach, on the other hand, combines the advantages of docking complete molecules with an assessment of synthesizability by fragmenting the molecules and matching the resulting fragments against a database of building blocks. As long as no reaction mechanisms are provided, this does not guarantee synthesizability. However, it increases the likelihood sufficiently to impose meaningful constraints on the training data and the output of the molecular generative model without excessively restricting its ability to generate novel chemistry.

Despite these promising results, the approach has inherent limitations. The reliance on molecular docking as a proxy for experimental affinity introduces potential inaccuracies, as these binding potential predictions usually do not capture entropic effects or receptor flexibility. Additionally, while the building-block-based generative search increases the chances of the availability of precursors with matching molecular substructures, it does not guarantee the synthetic feasibility of the end products. However, the presented framework provides a foundation for further extensions. Especially coupling this evolutionary–AI optimization loop with retrosynthetic analysis tools [44, 45] could further improve the prediction of synthetic accessibility. Additionally, incorporating receptor ensemble docking [46] or molecular dynamics-based scoring [47] could further account for receptor flexibility and dynamic binding effects, potentially increasing the experimental hit rate of the virtual hits in the pipeline. Finally, adding experimental validation cycles into the loop could establish an autonomous, closed-loop discovery platform that iteratively learns from both computational predictions and wet-lab results, ultimately selecting the best lead candidates for a given drug target from ever-increasing virtual chemical compound libraries.

AI-driven evolutionary frameworks, such as the one presented here, may become the new benchmark for early-stage discovery campaigns and potentially supersede HTS, which largely remains the current gold standard for early-stage hit and lead discovery in the biotech and pharmaceutical industry to date. By enabling *in silico* exploration of accessible chemical space orders of magnitude larger than physical HTS libraries, such approaches hold the potential to identify hits with higher initial activity, improved selectivity, more favorable pharmacological properties, and reduced off-target liabilities. These features could not only accelerate the early phases of drug discovery but also help mitigate downstream risks by lowering the likelihood of costly late-stage failures in drug development.

## Methods

### Molecule generation

For molecule generation in each iteration, we used the previously published REINVENT framework, i.e. the REINVENT randomized implementation by Arous-Pous et al. [22, 23]. For generating a pretrained model (prior), we used the referenced dataset of ∼1.5 million ligands from the ChEMBL database. For pretraining, we followed the recommendations of the authors. In brief, the dataset was split into a training and validation set at a ratio of 10:1. Then, the SMILES were randomized 10-fold using the randomize function of the REINVENT randomized implementation. Model pretraining was performed for 100 epochs on an NVIDIA Tesla A100 GPU with an adaptive learning rate. The resulting prior model was used to sample molecules in the first iteration (the starting population). After each iteration, the top 20% plus a random set of an additional 4% of molecules were extracted from the current total ranking, split into a training and validation set (ratio 10:1), SMILES were randomized again by 10-fold, and the current model was further fine-tuned for 10 epochs with an adaptive learning rate. The resulting fine-tuned model was then used in the subsequent iteration to generate the next set of ligands.

### Molecule decomposition and fragment search

To prioritize molecules that can potentially be generated by the combination of Enamine REAL building blocks, each AI-generated molecule was decomposed into fragments using RDKit [48, 49]-based molecule decomposition based on the BRICS reaction rules [24]. The resulting fragments were then compared to a database of molecular building blocks (∼1 million) from Enamine that was provided by Chemspace Ltd. Only molecules for which each fragment could be found in the building block database were further considered for virtual screening. For fast database searches of the fragments of each generated molecule, we used Morgan fingerprint [50]-based Tanimoto similarity [51] searching with a cutoff of 1.0 (exact match) using GPUsimilarity [52].

### Target and ligand preparation

As a target protein, we used the previously resolved x-ray crystallographic structure of the human MOR in complex with fentanyl (PDB: 8EF5) [26]. The protein structure model was cropped (amino acids 66–351), stripped of small molecule compounds, and side chain protonation states were generated using PROPKA 3 [53, 54] and PDB2PQR [55, 56]. The structure was prepared with side chain protonation states at pH 7.4 and 6.5, respectively. The resulting structures (PDB files) were imported into AutoDock-Tools (ADT) [57]; non-polar hydrogens were removed, and a PDBQT file was exported. For ligand preparation, we used the VirtualFlow for Ligand Preparation (VFLP) module of VirtualFlow [8, 21]. As settings, each generated molecule (SMILES) was first desalted, neutralized, and then the dominant protonation state at pH 7.4 or 6.5 was generated, respectively. For p*K_a_* predictions of the ionizable groups of the generated molecules, we used the hybrid ML- and semiempirical quantum mechanics (QM)-based method QupKake [29]. RDKit was then used to generate the final protonation states based on the p*K_a_* prediction. From each pH-specific molecule, a three-dimensional conformer was generated using Molconvert from Chemaxon. Finally, using Open Babel [58, 59] a PDBQT ligand file was generated for molecular docking.

### Molecular docking

For high-throughput molecular docking, we used VirtualFlow for Virtual Screening (VFVS), which is an open-source platform for ultra-large virtual screening on high-performance computing (HPC) clusters and in the cloud [8]. The target area (docking box) was centered onto the bound fentanyl (PDB: 8EF5) [26] and the area was selected to be 15 × 15 × 15 Å in size. As a docking program, we used Quick Vina 2 [60] with an exhaustiveness of 3, and each ligand was docked in three independent docking runs (*n*=3).

### Experimental validation using radioligand binding assay

Human embryonic kidney 293 (HEK293) cells stably expressing rat MOR were cultured in T175 flasks and harvested by washing with ice-cold 50 mM Tris-HCl (pH 7.4), followed by scraping, homogenization, and centrifugation at 42,000×g for 20 min at 4*^◦^C*. The resulting membrane pellets were resuspended in assay buffer, stored at −80*^◦^C*, and prepared as described previously [25, 30]. Protein concentration was determined using the Bradford assay. Radioligand displacement assays were performed with [*H]-DAMGO (4 nM; ∼50 Ci/mmol) as a selective MOR tracer at defined extracellular pH values (7.4 and 6.5). Test compounds were dissolved in dimethyl sulfoxide (DMSO) and diluted in ice-cold 50 mM Tris-HCl buffer so that all samples, including vehicle controls, contained 1% (v/v) DMSO. This ensured compound solubility and consistent solvent conditions for normalization. Membranes were incubated for 90 min at room temperature with [*H]-DAMGO and increasing concentrations of test compounds (1 × 10^−8^ − 5 × 10^−5^ M, final). After incubation, samples were rapidly filtered under vacuum through polyethylenimine-treated GF/B glass-fiber filters, washed with ice-cold assay buffer adjusted to the respective pH, and the retained radioactivity was quantified by liquid scintillation counting (Tri-Carb 4910TR, PerkinElmer) after overnight extraction. For each experiment, specific [*H]-DAMGO binding in vehicle controls (1% DMSO, no competitor) was set to 100%, and binding at each concentration was expressed as a percentage of this control. Nonspecific binding was defined as binding in the presence of naloxone (10 µM) and was subtracted from total binding to obtain specific binding. All binding experiments and analyzes were conducted under blinded conditions. Normalized displacement data (% specific binding remaining vs. log [compound]) were fitted using nonlinear regression with a one-site competition model (GraphPad Prism) to determine the half maximal inhibitory concentration (IC_50_) values. Concentration series replicates were removed from the analysis if the nonlinear regression returned an R^2^ *<* 0.8, a Span *<* 5 or *>* 110, or if the fit was flagged as ambiguous, indicating non-unique parameter estimation. The IC_50_ values were calculated from a minimum of 3 independent membrane preparations (biological replicates) per condition and reported as mean ± SEM (or 95% CI). Statistical comparisons between pH conditions were made using unpaired, two-tailed t tests.

### Molecular dynamics simulations and contact analysis

To evaluate the plausibility of the binding mode and to further strengthen the hypothesis that the positively charged hydrogen atom of compound 1, under acidic conditions, mostly contributes to its pH-specific binding, we additionally performed MD simulations and trajectory analyzes. For setting up the simulation system, we exported the binding poses of the top scoring compounds for further analysis using MD. The simulation system was constructed with explicit membrane and solvent modeling. The MOR structural model (PDB: 8EF5) [26] along with the docked ligand was embedded within a phosphatidylethanolamine (POPE) lipid bilayer and equilibrated at 25 °C. The embedding was carried out using a reference structure of the MOR obtained from the OPM (orientations of proteins in membrane database) server [61, 62] to ensure appropriate spatial insertion. The orthorhombic membrane box had an 8 x 8 x 8 nm buffer around the protein that ensured adequate separation between the protein and the boundary. To maintain charge neutrality, chloride counterions were added as required. The ionic strength of the system was set to 0.15 M NaCl to mimic physiological salt conditions. All atoms were parameterized using the OPLS2005 force field [63] with TIP3P as the choice for water. The system was then minimized with restraints on heavy atoms and subjected to 100 ps of minimization with a small maximum displacement (*δ*_max_ = 1 Å) without positional restraints to remove steric clashes in the solvated membrane embedded system before thermal equilibration. For production runs, MD simulations were performed using the Desmond academic simulation package [64]. The production MD was carried out under NPAT ensemble conditions (constant number of particles, pressure, area, and temperature) [65] at 310.10 K to mimic physiological temperature. A total simulation time of 1 µs was sampled using the Martyna-Tobias-Klein barostat [66], the Nosé-Hoover chain thermostat [67, 68], and the RESPA multigrator [69]. The specialty of the RESPA integrator is the ability to separate short range and long range forces, or quickly changing and slowly changing forces, into different groups. The faster, short ranged forces are calculated three times (2 fs) as much as the slower, long range forces (6 fs). The non-bonded interactions were set to a cutoff radius of 9.0 Å. Initial velocities were assigned from a Maxwell-Boltzmann distribution at 310.10 K. No enhanced sampling or biasing protocols were utilized in this production phase. Each simulation setup was replicated with different initial random velocities at least three times. After MD simulations, the trajectories were processed for contact analysis. The vicinity cut-off was set to 6 Å. Protein residues with heavy atoms within 6 Å were retained for further analysis in each frame. Once the residue is in range, the interaction is flagged only after the specific atoms or centroids pertaining to the specific interactions, such as hydrophobic contacts, *π*-*π* stacking, or salt bridges. We used the ProLIF package for this purpose [70]. Down-stream analyzes focused on four residues (Ile146, Asp149, Met153, Tyr328) that we considered important based on the predicted binding pose.

## Code availability

Source code for the ligand preparation and molecular docking pipelines is available via the VirtualFlow Project [71] and on GitHub: VFLP and VFVS. The code of the molecular generative model developed by Arús-Pous et al. [22, 72] and the ChEMBL data set used for pretraining the model are also available on GitHub [23]. QupKake [29] was used for micro-p*K_a_* predictions, for which the source code is available on GitHub [73]. To generate protonation states based on QupKake’s results, we have used an in-house generated Python script using RDKit. For decomposition of AI-generated molecules and database fragment searching, we used RDKit and gpusimilarity from Schrödinger.

## Data availability

Molecular dynamics data trajectories (3 replicas each at pH 6 and 7.4), which also contain protein and ligand RMSD, RMSF, and Jupyter notebooks for analysis, are archived on Zenodo (DOI: *pending*) and securely stored internally at the Zuse Institute Berlin; they can be made available upon reasonable request.

## Acknowledgements

The authors gratefully acknowledge the computing time made available to them on the high-performance computer “Lise” at the NHR center NHR@ZIB. This center is jointly supported by the Federal Ministry of Education and Research and the state governments participating in the NHR (www.nhr-verein.de). The authors also thank Chemaxon for a free academic license.

