## Supplementary Information for "Evolutionary exploration of drug-like chemical space utilizing generative AI and virtual screening"

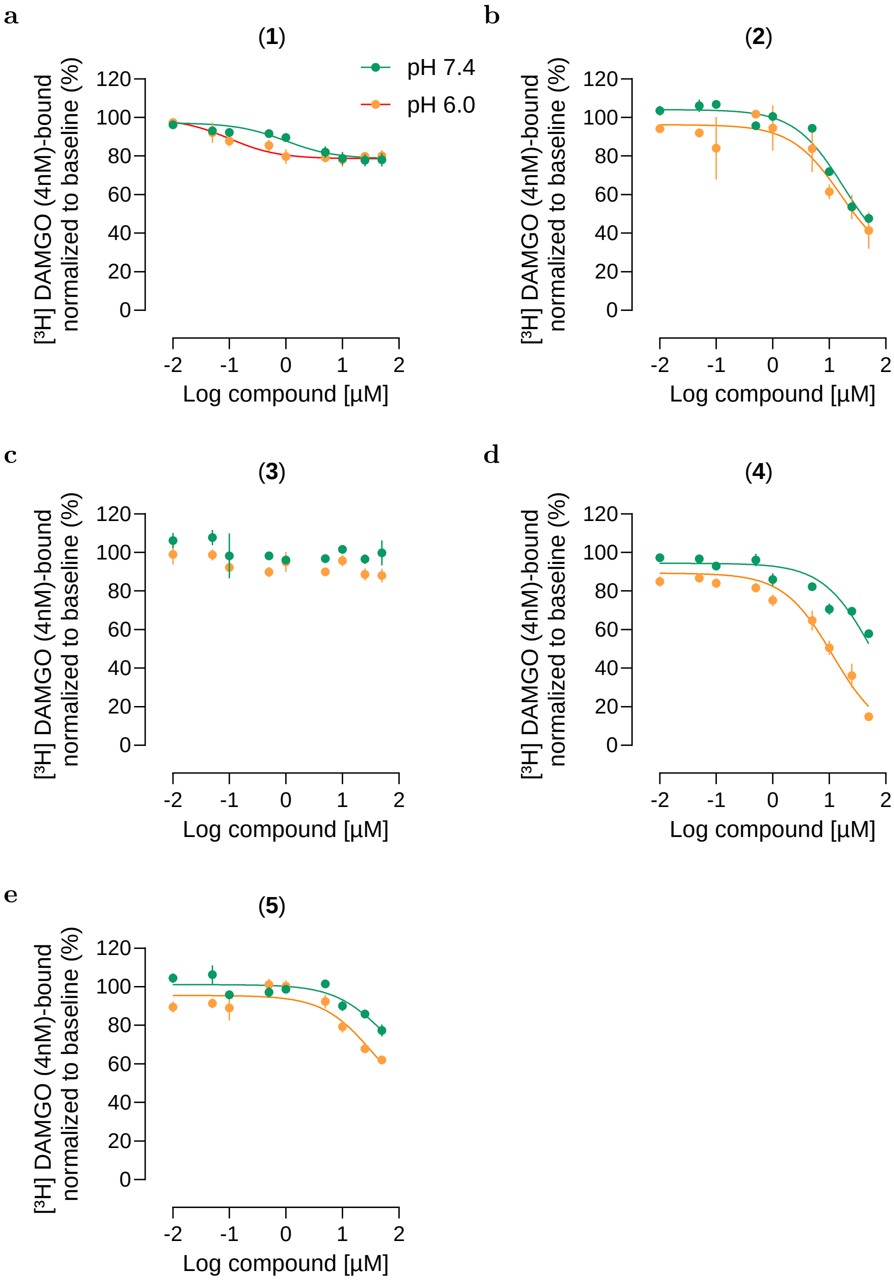


**Supplementary Fig. 1**. **Experimental validation of virtual hits.** (**a-e**) Radiolabeled RDA adding 10 nM to 50 µM of the selected hit compounds. Data points represent mean ± SEM.
